# µCT Scanning Effects on DNA and a Multi-Step Workflow for Archaeological Petrous Bones

**DOI:** 10.1101/2025.10.02.680114

**Authors:** Lumila Paula Menéndez, Pierre Luisi, María Clara López-Sosa, Sergio Monteiro Da Silva, Laura T. Buck, Susan Kuzminsky, Mirsha Quinto Sanchez, Anne Le Maitre, Christine Chappard, Cassandra Rios, Hans Groh, Wara Siles, Gustavo Montiel Hernández, Lorena Becerra-Valdivia, Juan Manuel Argüelles, Bernardo Yañez, Constanza de la Fuente Castro, Camila Tamburrini, Vivette Garcia-Deister, Miguel Angel Contreras Sieck, Sebastian Pasqualini, Paula Novellino, Candela Acosta Morano, Daniela Guevara, Daniela Mansegosa, Horacio Chiavazza, Sebastian Giannotti, Luis Tissera, Sebastián Pastor, Ivan Díaz, María Solange Grimoldi, Verónica Silva-Pinto, Ana Solari, Anne-Marie Pessis, Ramiro Barberena, Nicolás Rascovan

**Affiliations:** Escuela de Antropología, Universidad de Costa Rica, San José, Costa Rica; Department of Evolutionary Biology, University of Vienna, Vienna, Austria; Instituto de Antropología de Córdoba (IDACOR), Consejo Nacional de Investigaciones Científicas y Técnicas (CONICET), Córdoba, Argentina; Departamento de Antropología, Universidad Nacional de Córdoba, Córdoba, Argentina; Programa de Referencia y Biobanco Genómico de la Población Argentina (PoblAr), Argentina; Institut Pasteur, Université de Paris Cité, CNRS UMR 2000, Microbial Paleogenomics Unit, Pais, France; Department Anthropology of the Americas, University of Bonn, Bonn, Germany; Universidade Federal de Pernambuco, Recife, Brazil; Research Centre for Evolutionary Anthropology and Palaeoecology, School of Biological and Environmental Sciences, Liverpool John Moores University, Liverpool, United Kingdom; Department of Anthropology, California State University Channel Islands, Camarillo, United States of America; Instituto de investigaciones Antropológicas, Universidad Nacional Autónoma de México, Mexico City, Mexico; Konrad Lorenz Institute for Evolutionary and Cognition Research, Klosterneuburg, Austria; Laboratoire d’Imagerie Biomédicale, Sorbonne University, Paris, France; Universidad Nacional de La Plata, La Plata, Argentina; Laboratorio de Tecnologías Aditivas, Universidad Mayor de San Andrés, La Paz, Bolivia; Coordinación Nacional de Antropología, Instituto Nacional de Antropología e Historia, Mexico City, Mexico; Department of Anthropology and Archaeology, University of Bristol, Bristol, United Kingdom; Linacre College, University of Oxford, Oxford, United Kingdom; Dirección de Antropología Física, Instituto Nacional de Antropología e Historia, Mexico City, Mexico; Instituto de Ciencias Biomédicas, Facultad de Medicina, Universidad de Chile, Santiago de Chile, Chile; Centro de Investigación sobre el Envejecimiento, Cinvestav-Sede Sur, Mexico City, Mexico; Facultad de Ciencias, Universidad Nacional Autónoma de México, Mexico City, Mexico; Department of Anthropology, University of Minnesota, Minnesota and St. Paul, United States of America; Instituto Nacional de Antropología y Pensamiento Latinoamericano (INAPL), Consejo Nacional de Investigaciones Científicas y Tecnológicas (CONICET), Buenos Aires, Argentina; Consejo Nacional de Investigaciones Científicas y Tecnológicas (CONICET), Buenos Aires, Argentina; Museo de Ciencias Naturales y Antropológicas Juan Cornelio Moyano, Mendoza, Argentina; Instituto Interdisciplinario de Ciencias Básicas (ICB), Consejo Nacional de Investigaciones Científicas y Tecnológicas (CONICET), Universidad Nacional de Cuyo, Mendoza, Argentina; Laboratorio de Arqueología Histórica, Instituto de Arqueología y Etnología, Facultad de Filosofía y Letras, Universidad Nacional de Cuyo, Mendoza, Argentina; Museo Arqueológico Cerro Colorado, Agencia Córdoba Cultura, Cerro Colorado, Argentina; Instituto Regional de Estudios Socioculturales (IRES), Consejo Nacional de Investigaciones Científicas y Tecnológicas (CONICET), San Fernando del Valle de Catamarca, Argentina; Facultad de Filosofía y Letras, Universidad de Buenos Aires, Buenos Aires, Argentina; Universidad Isalud, Buenos Aires, Argentina; Instituto de las Culturas (IDECU), Consejo Nacional de Investigaciones Científicas y Técnicas (CONICET), Buenos Aires, Argentina; Programa de Doctorado en Geografía e Historia del Mediterráneo desde la Prehistoria a la Edad Moderna, Universitat de València, Valencia, Spain; Facultad de Ciencias Sociales e Historia, Universidad Diego Portales, Santiago de Chile, Chile; Área de Antropología, Museo Nacional de Historia Natural, Santiago de Chile, Chile; Fundação Museu do Homem Americano (FUMDHAM), Sao Raimundo Nonato, Brazil; Centro de Investigación, Innovación y Creación (CIIC-UCT), Facultad de Ciencias Sociales y Humanidades, Universidad Católica de Temuco, Temuco, Chile

**Keywords:** petrous bone, ancient DNA preservation, computed tomography, digital preservation, sustainable and interdisciplinary research, ethics in biological research

## Abstract

The petrous portion of the temporal bone is a key element in human evolutionary studies due to its exceptional preservation of biomolecules and morphological information. However, intensive and often redundant sampling has raised concerns about sustainability and long-term conservation. Here we present the first systematic evaluation of whether micro-Computed Tomography (µCT)—a widely used tool for digital preservation—affects ancient DNA (aDNA) integrity in human petrous bones. We analyzed 93 archaeological samples from Argentina, of which 50 had been scanned using µCT and 43 had not. We compared six molecular parameters, including endogenous content, read length, cytosine deamination patterns and contamination estimates. No statistically significant differences were observed between scanned and unscanned samples across any parameter (Mann-Whitney/Wilcoxon tests, p <0.05). Although mitochondrial contamination was marginally higher in scanned samples (p = 0.051), this was not driven by contamination estimates above the widely accepted 5% threshold for genomic analysis, Moreover, this pattern was not observed when considering nuclear contamination. These results indicate that, under appropriate scanning conditions, µCT imaging does not compromise DNA preservation. Building on this evidence, we propose a sustainable, multi-step workflow that integrates biological profiling, osteobiography, imaging, and compositional pre-screening prior to molecular sampling. This interdisciplinary approach maximizes the scientific information obtained from skeletal collections while minimizing destructive practices, thereby promoting ethical and sustainable research on irreplaceable anthropological remains, and fosters collaboration across research fields.

## 1. Background

Over the past two decades, advances in archaeometry—including imaging techniques and next-generation DNA sequencing—have significantly enhanced our understanding of human evolutionary history (Weber & Bookstein, 2011; Pääbo, 2014). In particular, the search for optimal sources of ancient DNA (aDNA) has led to the widespread adoption of petrous bone sampling, due to its exceptional biomolecular preservation (Pinhasi et al., 2015, Hansen et al., 2017; Pilli et al., 2018; Pinhasi et al., 2019). Initially, teeth were considered the best skeletal source of aDNA because of their relative resistance to contamination (Rohland & Hofreiter, 2007; Higgins & Austin, 2013; Kendall et al., 2018). However, recent research has shown that the cochlea within the petrous portion of the temporal bone (Figure 1a-d) exhibits even better DNA preservation, outperforming all other anatomical structures (Pinhasi et al., 2015; Hansen et al., 2017; Pilli et al., 2018; Gaudio et al., 2019; Pinhasi et al., 2019). More recently, despite the considerably lower preservation of ear ossicles (Figure 1d) in archaeological samples, they have been shown to yield DNA of comparable quality and complexity to that of the cochlea, leading to the recovery of similar amounts of endogenous DNA, mtDNA coverage, nuclear SNP coverage, and number of SNPs called (Sirak et al., 2020). Other studies focused on preserved ancient molecules—such as stable isotopes and radiocarbon dating— have followed this trend, favoring the petrous bone for sampling (e.g., Jørkov et al., 2009; Douka et al., 2017; Price et al., 2018). As a result, the petrous bone (Figure 1c) has become the preferred target for most molecular studies, being targeted and primarily used for DNA analysis, and consequently employed in radiocarbon dating, ZooMS, and stable isotope studies when multiple analyses are conducted on the same individual or sample.

**Figure 1.**
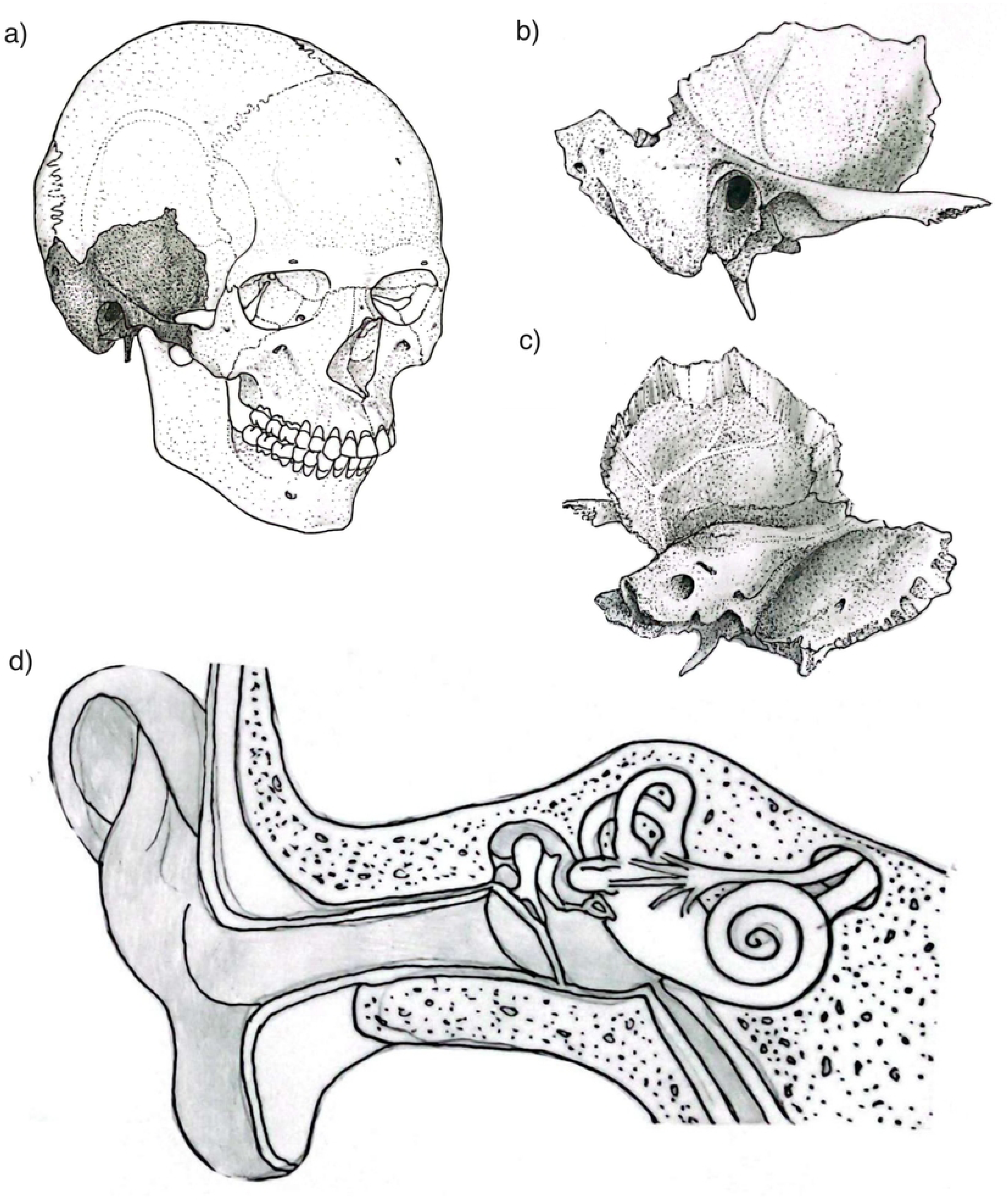
Main anatomical features of the temporal bone and petrous portion: a) Human skull with the temporal bone highlighted, b) temporal bone from lateral view, c) temporal bone from internal view showing the petrous portion; d) internal detail of the petrous bone showing the location and position of the inner and middle ear. The drawings were created by Sergio Monteiro Da Silva.

Beyond its molecular significance, the petrous bone also holds considerable value for morphological and other evolutionary studies. The cranial base and temporal bone have long been recognized as cranial regions that are less susceptible to environmental influences and preserve a strong phylogenetic signal, which are useful for reconstructing population histories (Harvati & Weaver, 2006; Collard & Wood, 2000; Smith, 2009; von Cramon-Taubadel, 2009; Reyes-Centeno et al., 2017). More recently, the bony labyrinth within the petrous portion (Figure 1d) has been identified as an anatomical structure that reliably reflects random evolutionary changes and, as such, retains both phylogenetic and population-level signals (Le Maître et al., 2017; Ponce de León et al., 2018; Uhl et al., 2022; Smith et al., 2024; Urciuoli et al., 2025). As a result, the petrous bone has become a key resource across multiple research domains.

The growing demand for petrous bone samples, however, has not been accompanied by a sustainable research management strategy. Despite attempts to reduce damage and implement minimally invasive protocols, current practices often still lead to the destruction of the cochlea, preventing detailed analysis of the bony labyrinth, and impeding the study of temporal and basicranial forms (Sirak et al., 2017). The rapid expansion of aDNA studies has led to what some researchers describe as “petrous fever”—a facet of the broader “aDNA rush” or “aDNA frenzy”. This trend is mostly characterized by extensive sampling practices that underscores preservation arguments and ethical recommendations, particularly by large laboratories. These labs often operate without clearly defined research questions and instead pursue exploratory objectives, raising concerns about the long-term sustainability of such practices (Sykes et al., 2019; Nieves-Colón et al., 2020; Argüelles et al., 2022; Källén et al., 2024; Menéndez et al., 2025). The emphasis on generating genetic data frequently comes at the expense of other valuable sources of biological information—such as morphology, histology, and isotopic analyses—which provide critical insights into past human health, diet, mobility, and cultural practices.

Yet, the momentum of the aDNA rush has not been accompanied by systematic efforts to digitally preserve samples prior to their destruction. One widely available approach is CT scanning, which uses X-rays and computer algorithms to generate detailed cross-sectional and 3D reconstructions of internal structures (Weber & Bookstein, 2011; Menéndez et al., 2024). Two main types are commonly used: medical CT, which captures entire human bodies at resolutions of 0.5–1 mm, and µCT, which offers much finer resolution (1–100 µm) for small samples and is especially useful for applications requiring detailed structural analysis, such as studies of the inner ear. Despite its benefits for digital preservation, CT scanning raises concerns regarding the potential effects of X-ray exposure on biomolecular preservation. Since CT uses ionising radiation, it may damage biological molecules, including DNA, and thereby compromise the integrity of ancient genetic material in bone (Immel et al., 2016).

Recent studies in both contemporary and prehistoric animals suggest that, when appropriate CT scanning protocols are applied, conventional μCT scanning does not reach radiation doses considered harmful for DNA (Paredes et al., 2012; Hall et al., 2015; Immel et al., 2016). Similar results have been reported for X-ray radiography of 18th-century and Bronze Age human femurs (Fehren-Schmitz et al., 2016). In such cases, DNA degradation does not exceed the natural rate of temporal decay. These findings support the safe use of radiography and μCT scanning in bioarchaeological studies, provided that suitable imaging practices are followed. Yet, it remains unclear whether these conclusions hold for archaeological human petrous bones—currently the most targeted element for aDNA retrieval—and under the specific imaging conditions of μCT scanning typically used in contemporary biological and evolutionary anthropology research.

Our study is the first to systematically address this question in archaeological petrous bones. Unlike animal samples or archaeological femora, the petrous is the densest human bone and the principal reservoir for ancient DNA, so its response to μCT radiation cannot be directly inferred from other anatomical elements. Given that petrous bones are now the primary source for paleogenomic research, even minor effects on DNA integrity may have disproportionately large implications compared to other skeletal elements. Human petrous bones also differ from animal samples in their microstructure, being far denser and less porous, which makes them a unique case that requires direct evaluation rather than extrapolation from other species.

In this paper, we advocate for an urgent shift in the research design and management of human remains for paleogenomics, bioarchaeology and human evolutionary studies. To support this call, we present the results of an experiment comparing DNA preservation in 93 petrous samples from archaeological sites in Argentina (the Andes, Central Argentina, and Cuyo), spanning the Middle to Late Holocene. Some of these samples underwent μCT scanning, while others from the same assemblages did not (Table S1). Building on these findings, we propose a sustainable workflow that maximizes the biological information derived from the petrous portion of the temporal bone, while also addressing the challenges posed by intensive sampling from skeletal collections worldwide. Although implementing such a strategy may present obstacles, we argue that this practice will ultimately advance scientific knowledge production. While our discussion is grounded in the field of human evolutionary research, its implications extend more broadly to forensic, paleontological, and biological sciences.

## 2. Relevance of the temporal bone and its petrous portion

The petrous portion is a dense, pyramidal component of the temporal bone—a thick, bilateral structure that forms part of the skull’s lateral walls and base. The temporal bone itself comprises several parts: the squamous, tympanic, styloid, and petrous portions (Reisser et al., 1996; Chae & Rodriguez Rubio, 2020; Figure 1a). Of these, the petrous portion is the most robust, housing critical auditory and vestibular structures.

The petrous portion is positioned between the sphenoid and occipital bones and divides the middle and posterior cranial fossae. It consists of three primary surfaces: anterior/superior, posterior, and inferior (Figure 1b-c). The anterior/superior surface forms part of the middle cranial fossa and extends from the arcuate eminence to the petrous apex. The posterior surface continues into the posterior cranial fossa and contains the internal auditory meatus, which serves as a passage for the facial (cranial nerve VII) and vestibulocochlear (cranial nerve VIII) nerves. The inferior surface houses multiple foramina that transmit essential cranial nerves and blood vessels (Ekdale, 2016; Lim & Brichta, 2016).

Internally, the petrous portion encloses several anatomical structures, with the inner and middle ear being the ones composed of bone (Figure 1d). The inner ear, or bony labyrinth, is a highly complex structure responsible for hearing and balance. It is located in the petrous portion of the temporal bone, near its center and oriented medially and slightly anteriorly (Figure 2). To illustrate its location, position, and orientation, we prepared a video showing an inner ear within a temporal bone analyzed in this work (Figure 2; López-Sosa et al., 2025). The inner ear consists of the cochlea, semicircular canals, and vestibule (Figure 1d). The cochlea is a spiral-shaped organ that converts mechanical sound waves into neural signals, which are then transmitted to the brain via the auditory nerve. The semicircular canals detect rotational movements of the head, which aids in balance and spatial orientation. The vestibule plays a key role in detecting linear acceleration and head positioning relative to gravity. The middle ear is an air-filled cavity that plays a crucial role in sound transmission. It is located between the tympanic membrane and the inner ear and contains the ossicles (malleus, incus, and stapes) (Figure 1d). Other non-bony structures contained within the petrous portion include the membranous labyrinth, internal acoustic meatus, carotid canal, jugular foramen, and facial canal. Additionally, two smaller bony structures are present: the arcuate eminence and the tegmen tympani (Ades et al., 1974).

**Figure 2.**
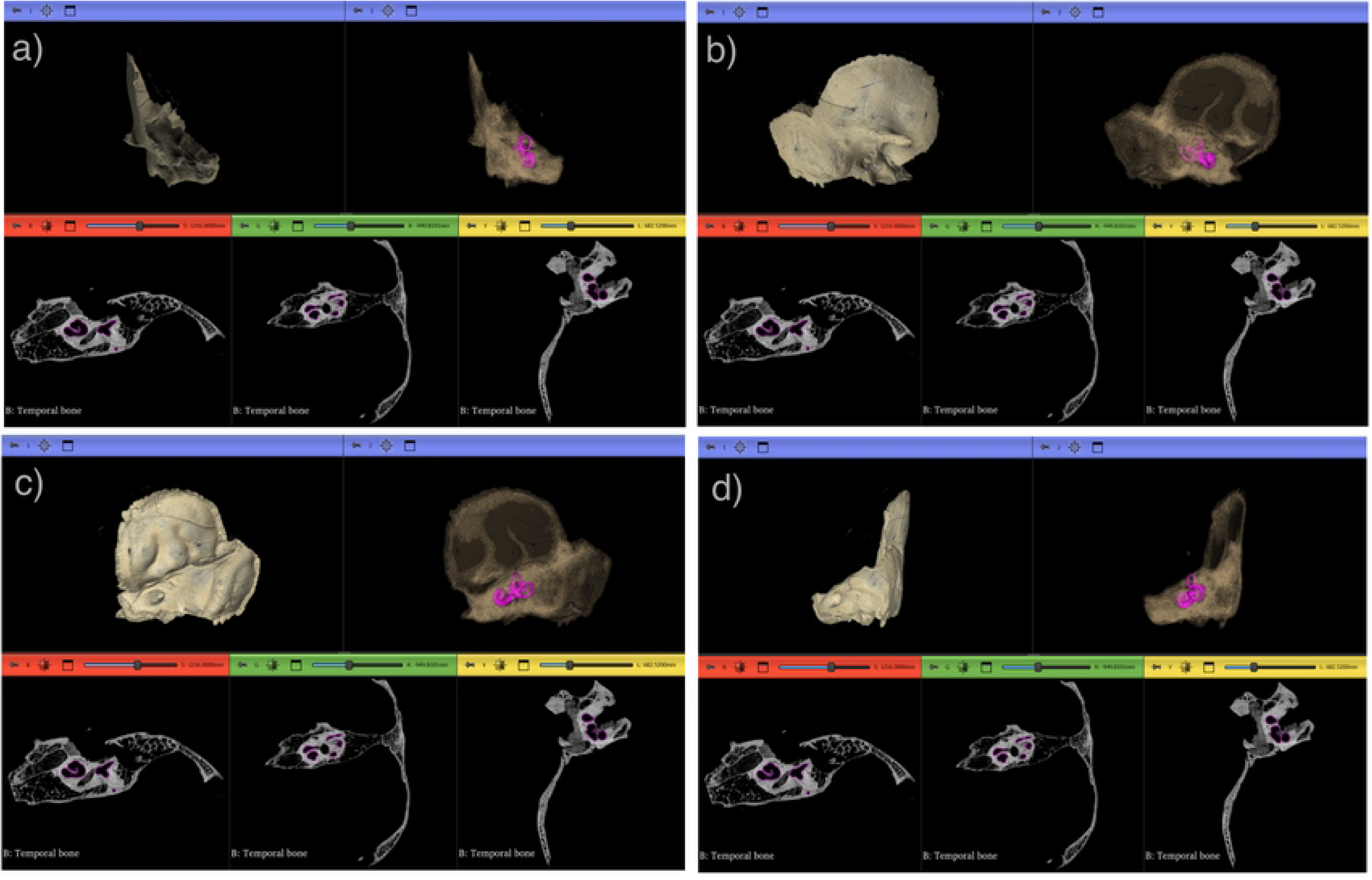
µCT scans of a temporal bone showing the segmented inner ear or bony labyrinth (highlighted in magenta) in a) anterior, b) lateral, and c) postero-lateral. In each image a-d, the two upper panels display 3D models of the temporal bone and the inner ear (the left one with texture, and the transparent right one showing the inner ear). The three lower panels show 2D slices in the anatomical orthogonal planes indicated above: R(Red) = axial, G(Green) = sagittal, and Y(Yellow) = coronal. This Figure was created using 3D Slicer Preview Release 5.9.0 (Fedorov et al., 2012; https://www.slicer.org) and Inkscape.

One of the most distinctive features of the petrous portion of the temporal bone is its early development and exceptional stability over time, making it highly valuable for research across multiple disciplines. Fully formed by the eighth week of intrauterine life, the petrous undergoes little to no postnatal remodeling during adulthood—unlike most other skeletal elements. It is also the densest and most highly mineralized part of the human skeleton (its name derived from the Latin *petrosus*, meaning “rock-like”), traits that contribute to its remarkable preservation in both archaeological and fossil contexts. These properties not only protect the bone from post-mortem degradation but also make it one of the most reliable sources for the recovery of ancient DNA. Its compact structure consistently yields high quantities of endogenous DNA, making it the most sought-after skeletal element in molecular analysis of ancient humans (Jeffery & Spoor, 2004; Parker et al., 2020; Ponce de León et al., 2018; Sirak et al., 2020). In addition to its utility for molecular analyses, the petrous bone’s structural stability also ensures the preservation of fine morphological details, which are often lost in other skeletal elements due to taphonomic processes. This makes it an equally valuable resource for studies focused on skeletal morphology.

## 3. A Proposal for a Sustainable Workflow in Petrous Bone Studies: Benefits and Challenges

As described above, the petrous portion of the temporal bone holds exceptional informational value across multiple disciplines, making careful research design essential when planning studies involving its sampling for molecular analysis. Given this significance, we issue an urgent call to optimize workflows that maximize the preservation of biological information from this structure. Implementing such measures is crucial for safeguarding data and ensuring its availability, not only for current researchers, but also for future generations of scientists (Fox & Hawks, 2019).

During visits to collections across different continents, some of us have encountered cases in which both petrous bones from a single individual were entirely removed—one for genetic analysis and the other for radiocarbon dating, or in some cases, both for separate attempts at DNA extraction, sometimes by different laboratories—even when a single sample would have likely sufficed (Squires et al., 2019). Yet it remains unclear how many of these samples were digitally preserved prior to destructive analysis—likely only a small fraction. This often depends on whether principal investigators secured funding or resources for digital preservation—which is rarely the case—or collaborated with morphologists who had both access to the necessary resources and a specific interest in studying this structure, as well as the foresight to recognize the value of digitally preserving the samples.

After almost a decade of intensive petrous bone sampling for genomic research, it is imperative to reassess current practices in light of long-term sustainability and responsible resource use. The petrous portion has become the preferred skeletal element for aDNA retrieval (Pinhasi et al., 2015; Hansen et al., 2017; Pilli et al., 2018; Pinhasi et al., 2019), contributing to the generation of over 10,000 ancient human genomes in the past decade (Mallick et al., 2024). In fact, according to the metadata associated with the curated compendium of ancient human genomes, the Allen Ancient DNA Resource (AADR v9; Mallick et al. 2024), aDNA data has been retrieved from petrous bones in 47% of all sequenced individuals. Importantly, the petrous bone was the only sampled bone in 96% of these cases. However, large-scale genetic data production often comes at the expense of other valuable lines of inquiry (Källén et al., 2024; Menéndez et al., 2025). Given the finite nature of skeletal collections, ethical considerations surrounding destructive sampling, and evolving sampling methods, it is crucial to adopt a more balanced, multi-proxy research strategy—one that ensures paleogenetic analyses complement rather than overshadow other research methods. By embracing such a framework, the field can move toward a more comprehensive and sustainable use of the petrous bone, preserving its scientific value while continuing to deepen our understanding of past populations.

Some researchers have implemented protocols to ensure the sustainable use of information contained in petrous bones. To this end, the same petrous bone is used for multiple purposes: before it is partly pulverized to generate molecular data (both genomic and isotopic), biological anthropologists perform CT and surface scans to capture its morphological information. While this sequence of operations may slow the production of genomic and isotopic data, we believe it enables researchers to fully harness the diverse information embedded in bone. Incorporating digitalization protocols prior to sampling for genomic and isotopic data, enhances research workflows and maximizes the biological information provided by multiple independent sources. Moreover, these practices foster interdisciplinary collaboration, support the integration of multiple lines of evidence on a single issue, and ensure the digital preservation of valuable morphological data for future research. In contrast, ongoing disciplinary fragmentation often results in researchers working in isolation, which can limit the potential for meaningful cross-disciplinary insights and comprehensive interpretations (Menéndez et al., 2022).

## 4. Testing the Effects of µCT Scanning on aDNA Preservation in a set of Petrous Fragments

Because CT scanning relies on X-ray radiation, we sought to evaluate whether such exposure could affect the authenticity, quality, and/or contamination levels of aDNA extracted from archaeological petrous bones that were µCT-scanned and compared to others from the same site that were not. To address this, we compared six key parameters commonly used in aDNA research: (1) endogenous DNA content; (2) average read length; frequency of cytosine-to-thymine (C→T) substitutions— analyzed separately for libraries that were UDG-treated (3) and those that were not (4); contamination estimates, which were based on mitochondrial DNA (mtDNA) using ContamMix (5), and (nuclear DNA contamination estimates using X chromosome-based methods for male individuals (6) (Table 1).

**Table 1.**
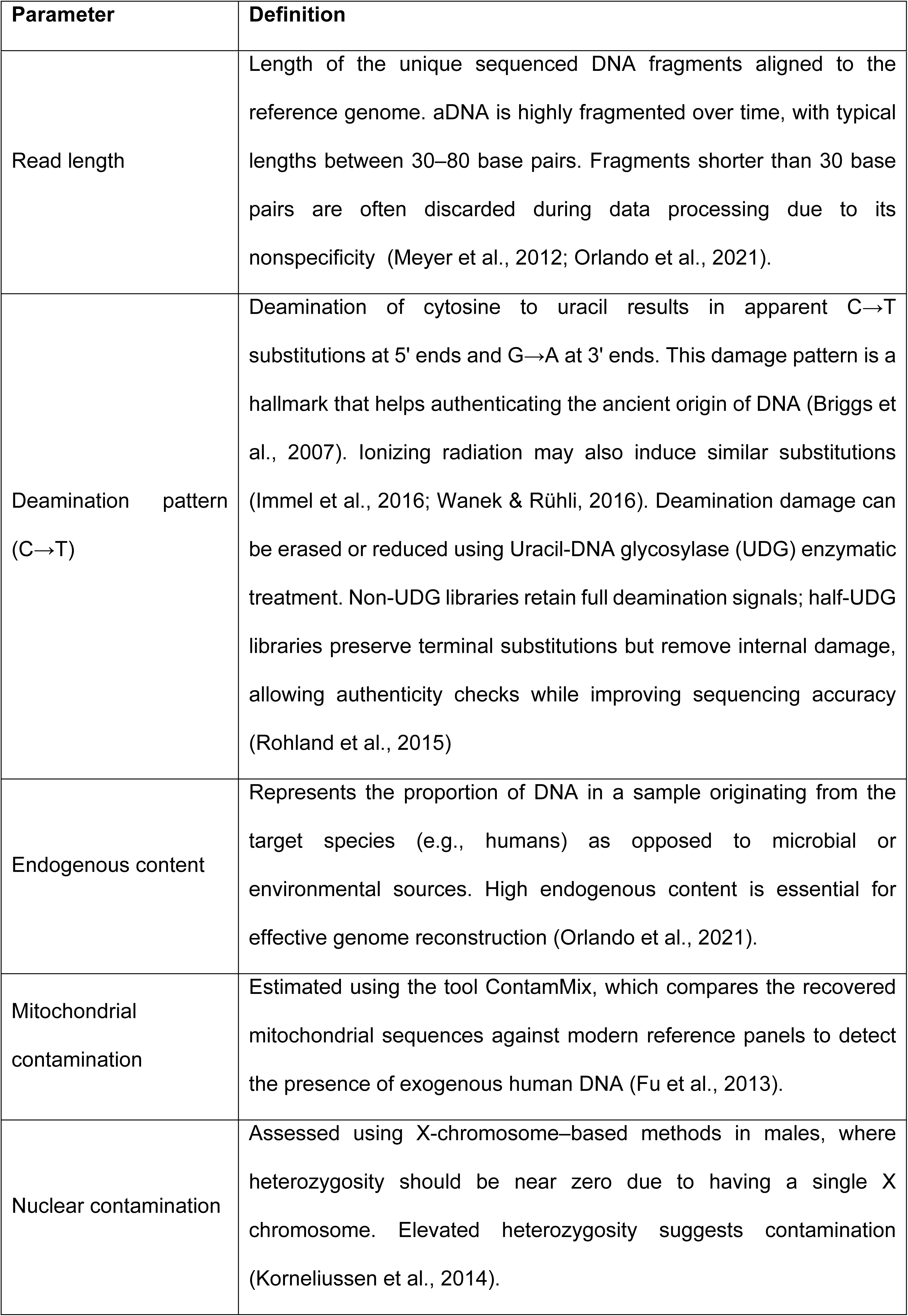
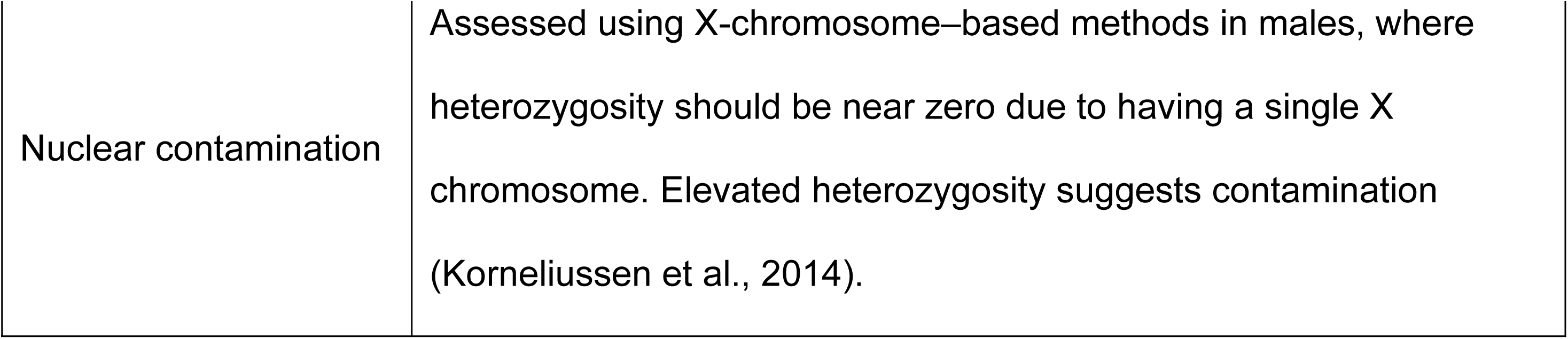
Key variables for assessing ancient DNA authenticity, quality, and contamination.

| Parameter | Definition |
| --- | --- |
| Read length | Length of the unique sequenced DNA fragments aligned to the reference genome. aDNA is highly fragmented over time, with typical lengths between 30–80 base pairs. Fragments shorter than 30 base pairs are often discarded during data processing due to its nonspecificity (Meyer et al., 2012; Orlando et al., 2021). |
| Deamination pattern (C→T) | Deamination of cytosine to uracil results in apparent C→T substitutions at 5' ends and G→A at 3' ends. This damage pattern is a hallmark that helps authenticating the ancient origin of DNA (Briggs et al., 2007). Ionizing radiation may also induce similar substitutions (Immel et al., 2016; Wanek & Rühli, 2016). Deamination damage can be erased or reduced using Uracil-DNA glycosylase (UDG) enzymatic treatment. Non-UDG libraries retain full deamination signals; half-UDG libraries preserve terminal substitutions but remove internal damage, allowing authenticity checks while improving sequencing accuracy (Rohland et al., 2015) |
| Endogenous content | Represents the proportion of DNA in a sample originating from the target species (e.g., humans) as opposed to microbial or environmental sources. High endogenous content is essential for effective genome reconstruction (Orlando et al., 2021). |
| Mitochondrial contamination | Estimated using the tool ContamMix, which compares the recovered mitochondrial sequences against modern reference panels to detect the presence of exogenous human DNA (Fu et al., 2013). |
| Nuclear contamination | Assessed using X-chromosome–based methods in males, where heterozygosity should be near zero due to having a single X chromosome. Elevated heterozygosity suggests contamination (Korneliussen et al., 2014). |
| Nuclear contamination | Assessed using X-chromosome–based methods in males, where heterozygosity should be near zero due to having a single X chromosome. Elevated heterozygosity suggests contamination (Korneliussen et al., 2014). |

Prior to scanning, petrous bones were kept inside their archival polyethylene bags, which limited direct handling and minimized the risk of contamination. Up to four samples were scanned simultaneously in the chamber under these conditions. Gloves and masks were worn during the placement and retrieval of the bags, and scanner components were cleaned regularly between runs to reduce potential cross-sample contamination.

The parameters used for µCT scanning are presented in Table 2. These values were selected to achieve sufficient resolution for morphological analyses while minimizing exposure, following recommendations from previous studies on radiological effects on biomolecules (Hall et al., 2015; Immel et al., 2016). The procedures followed for ancient DNA extraction, library preparation, and sequencing are the same as described in detail in Rascovan et al. (2025).

**Table 2.**
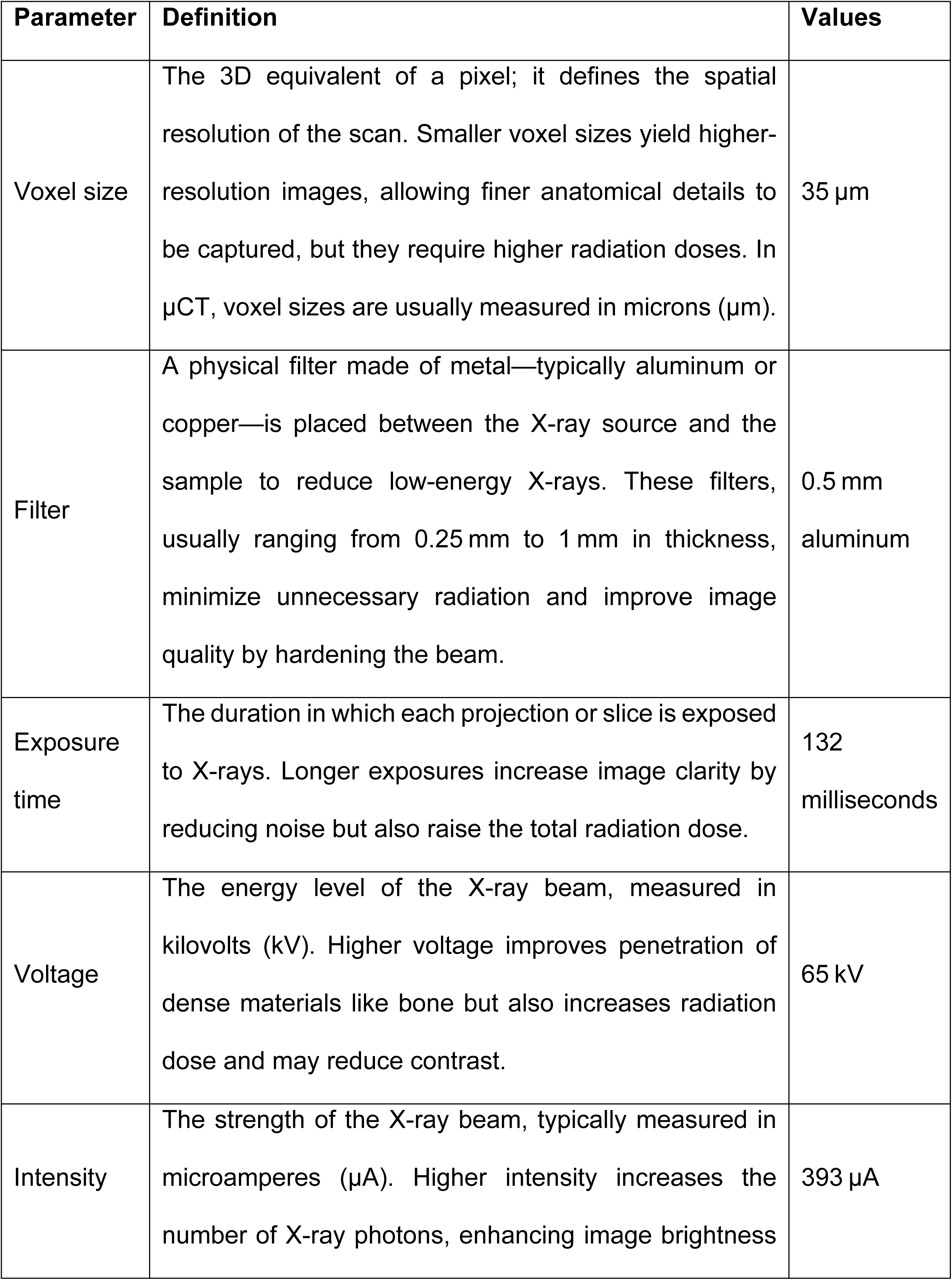
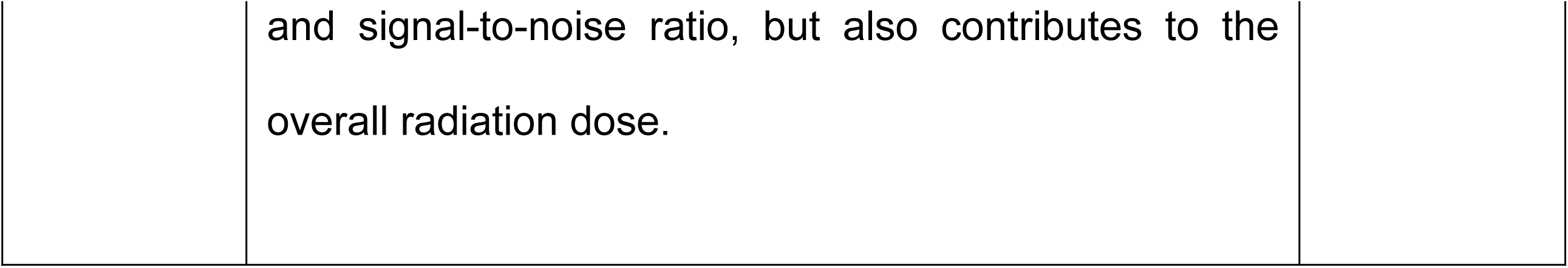
Description and operational values of µCT-Scanning parameters applied in the present study.

We analyzed petrous bone samples recovered from Middle and Late Holocene archaeological contexts across present-day Argentina (the Andes, Central Argentina, and Cuyo; Table S1), selecting samples that were either µCT-scanned (“True”) or not (“False”) prior to aDNA extraction (Figure 3). µCT scanning was conducted on 50 samples using a Skyscan 1176 Bruker scanner at the Faculty of Medicine, Paris Diderot University (Table 2). We sequenced aDNA libraries produced from these 50 scanned petrous samples and other 43 unscanned petrous that we used for our comparison. Although the samples came from similar archaeological contexts (i.e., same archaeological sites), they were not matched pairs (i.e., elements from the same individual), which limits our ability to attribute observed differences exclusively to µCT scanning.

**Figure 3.**
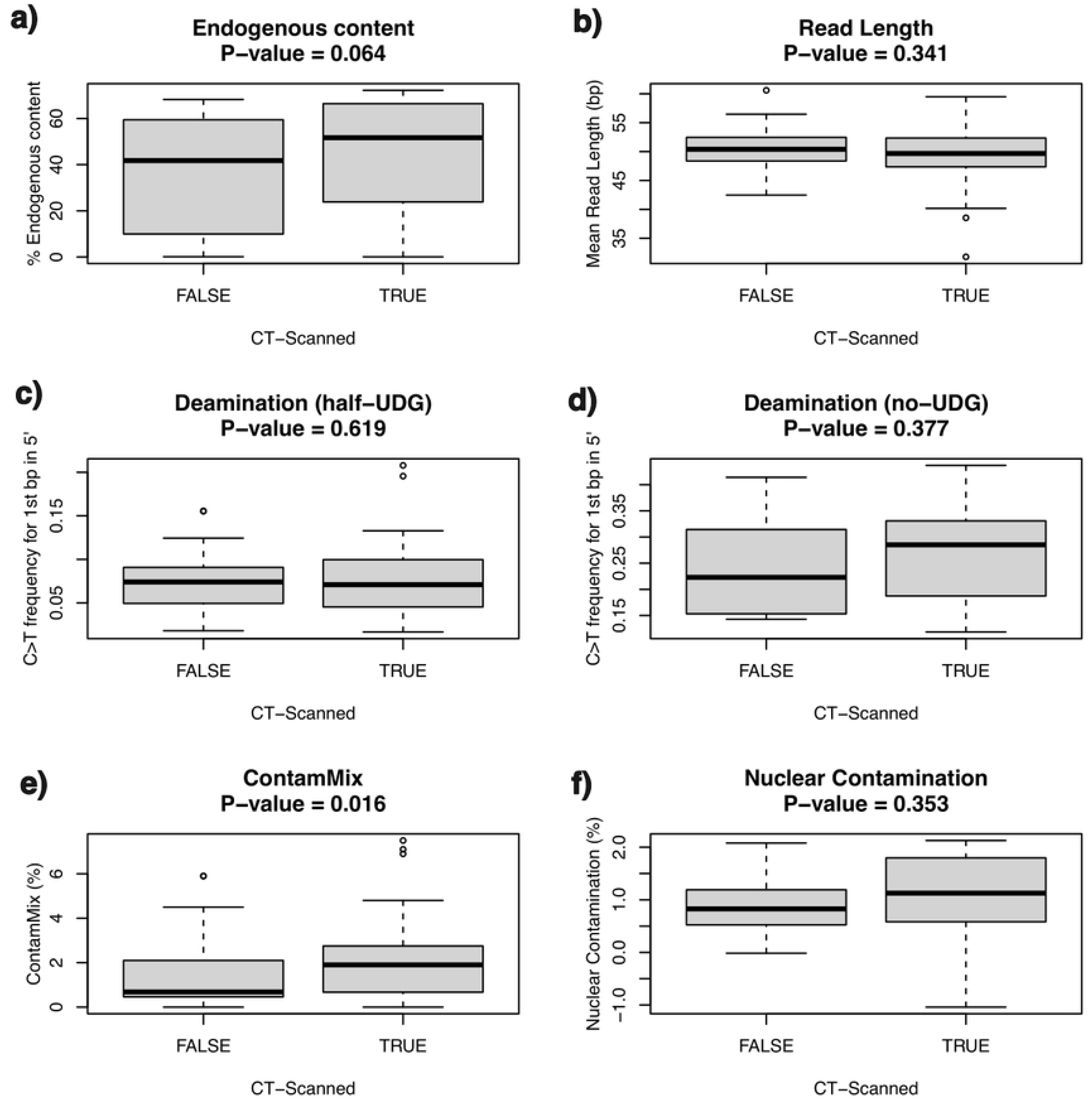
Comparison of aDNA preservation and contamination parameters in µCT-scanned (“True”; n = 50) and not µCT-scanned (“False”; n = 43) petrous samples: (a) Endogenous content: percentage of unique reads mapping to the human reference genome; (b) Average read length mapping to the human reference genome; (c) Frequency of C→T transitions at the 1st base-pair of the 5’ end of the reads for half-UDG treated DNA libraries, and for (d) non-UDG treated DNA libraries; Contamination estimates (in %) based on (e) mitochondrial DNA (with ContamMix, f) and nuclear DNA (f, only for male individuals). *P-*values are derived from a Wilcoxon rank-sum test conducted in R (R Core Team, 2024).

To assess potential differences between scanned and unscanned samples, we compared the distributions of the six parameters using non-parametric tests. Mann–Whitney U (Wilcoxon rank-sum) tests were carried out in R v4.3.2 (R Core Team, 2024), as the data did not follow a normal distribution. A threshold of p < 0.05 was used to evaluate significance. Because the samples were not matched pairs, this approach was considered appropriate for comparing independent groups. Descriptive statistics (medians and interquartile ranges) and parameter distributions were also examined to identify possible trends not captured by significance testing.

Table S1 in the Supporting Information provides a summary of the metadata for the analyzed individuals and DNA preservation results for each of the derived petrous samples. We did not find any statistically significant differences between the µCT-scanned and not µCT-scanned samples for any of the evaluated parameters (Mann Whitney U/Wilcoxon rank-sum test performed in *R*; p > 0.05) (Figure 3, Table 3). This suggests that, under the scanning conditions that we applied (Table 2), µCT imaging does not significantly compromise aDNA preservation, as assessed by the parameters considered here (Table 1).

**Table 3.** Wilcoxon rank-sum test results comparing µCT-scanned (n=50) and unscanned (n=43) petrous bones across key molecular parameters. No statistically significant differences were found (p > 0.05; see also Figure 3).

|  | U/W (Mann-Whitney/Wilcoxon statistic) | p-value |
| --- | --- | --- |
| Endogenous content | 1132 | 0.663 |
| Read length | 998 | 0.556 |
| Deamination (half-UDG) | 328 | 0.813 |
| Deamination (no-UDG) | 211 | 0.554 |
| ContaMix | 1301 | 0.051 |
| Nuclear contamination | 110 | 0.798 |

Read length and C→T substitution patterns in both half and non-UDG-treated libraries—commonly used indicators of aDNA fragmentation and authenticity—were similar between the two groups, providing no evidence of µCT-induced strand breaks or elevated damage. These findings align with previous studies suggesting that standard μCT protocols fall below radiation doses known to accelerate molecular degradation (Hall et al., 2015; Immel et al., 2016).

Endogenous content appeared to be very similar between the two groups (p = 0.663). Actually, the highest endogenous content was observed in the µCT-scanned samples. While this might seem counterintuitive, it likely reflects selection bias towards scanning better preserved bones rather than a biological effect of µCT scanning. In practice, researchers may prioritize samples that appear well-preserved, excluding more degraded specimens from imaging workflows.

Contamination estimates based on mitochondrial data were slightly higher in µCT-scanned samples (p = 0.051), although most (48 out of 49) remained below the commonly accepted 5% threshold for genomic analyses (Green et al., 2010; Fu et al., 2013). Although a marginal trend was detected in mitochondrial contamination, values remained below widely accepted thresholds and are best explained by sample selection rather than scanning effects. In contrast, no significant difference was observed between groups in nuclear contamination estimates (p = 0.798), with the highest value (2.3%) found in a sample that was not µCT-scanned.

In summary, our results provide no significant evidence that µCT scanning negatively impacts aDNA preservation, quality, or authenticity. Although minor trends were observed in some parameters, they are most parsimoniously explained by differences in sample selection or handling, rather than by the scanning process itself. In particular, the marginal increase in mitochondrial contamination (p = 0.051) is likely due to selection bias, as researchers may preferentially scan specimens that appear better preserved, rather than a biological effect of µCT scanning. These findings support the integration of µCT imaging into research workflows, if appropriate protocols are followed. By doing so, researchers can digitally preserve valuable morphological data prior to destructive sampling, and contribute to the broader goal of fostering a sustainable and interdisciplinary approach to human evolutionary research.

## 5. Proposed Workflow for a Sustainable Petrous Bone Research Agenda

To maximize the preservation of biological information, we propose that once access has been granted and the necessary permits to study the samples have been approved, specimens undergo a series of preliminary assessments before bone powder or fragments are collected for molecular analyses. We suggest following the recommendations of Prendergast and Sawchuk (2018) and Llamas et al. (2016), who offer good practice guidelines for handling human remains (e.g., wearing gloves, using sterile packaging). We do not suggest that the workflow proposed here should be the full responsibility of a single individual; rather, we envision this process as a collaborative and interdisciplinary team effort.

The multi-step workflow we suggest (Figure 4) is as follows:

1. Macroscopic evaluation: A detailed macroscopic assessment of bone preservation, diagenesis, taphonomy, and overall sample quality is key for making informed decisions about subsequent analyses (Price et al., 1992; Keenan, 2016).
2. Biological profiling and osteobiography analysis: The first step should be a comprehensive preliminary report that provides fundamental information about the skeletal elements of each individual. This includes estimations of sex and age at death, as well as the identification of notable pathological conditions such as trauma, disease, or enthesopathies, which may open new avenues for further investigation (Hosek & Robb, 2019). Once this foundational analysis is complete, an osteobiographical assessment can be undertaken. Osteobiography involves reconstructing an individual’s life history through the study of their skeletal remains, integrating biological data with archaeological and cultural context (Sofaer, 2006).
3. Digital preservation: Either medical, or preferably, µCT scanning—complemented by surface scanning or photogrammetry—should be conducted to create permanent digital records of samples. CT scanning enables the study of internal structures, while surface scanning and photogrammetry provide complementary data by preserving surface texture information. Together, these methods complement each other effectively (Weber & Bookstein, 2011; Menéndez et al., 2024). Recommendations for optimizing medical and μCT scanning parameters while minimizing DNA damage in dry bones and mummified remains can be found in Loynes and Bianucci (2021), Martin-Champetier et al. (2025), and Immel et al. (2016). To prevent redundant scanning, researchers should avoid repeating scans when comparable data are already available (Immel et al., 2016). In this context, data sharing—under the responsibility of national and private institutions and their curators— and the use of public or institutional repository databases play a crucial role in promoting efficient and sustainable research practices.
4. Bone pre-screening techniques to assess molecular preservation: In bone-collagen radiocarbon dating and stable isotope analysis, quick and cost-effective pre-screening techniques help assess collagen preservation and guide sampling strategies by identifying morphologically insignificant bone fragments that meet the thresholds for more labour-intensive and costly downstream analyses. Common approaches include measuring nitrogen content (%N) as a proxy for collagen presence (Brock et al., 2010), near-infrared spectroscopy (NIR), a non-destructive technique (Sponheimer et al., 2019), and X-ray fluorescence (XRF) analysis (Schakley, 2020). Zooarchaeology by mass spectrometry (ZooMS)—a relatively inexpensive, high-throughput technique—has also proven valuable in this context, particularly when integrated into species identification workflows (Harvey et al., 2016). Given the established correlation between collagen preservation and DNA survival (Kontopoulos et al., 2020; Sosa et al., 2013; Götherström et al., 2002), integrating these pre-screening techniques—alongside emerging minimally-destructive workflows—into multi-method approaches that incorporate aDNA analysis could enhance efficiency, reduce costs, and minimise destruction. Such strategies may also reduce reliance on sampling morphologically significant elements, such as the petrous bone, at the outset of molecular investigations.
5. Sampling for molecular analyses: Sampling should only proceed once steps 1–4 have been completed to ensure the most informed and responsible selection of samples. By conducting osteobiographical analysis, macroscopic evaluation, digital preservation, and XRF analysis beforehand, researchers can minimize unnecessary destruction and maximize the scientific value extracted from each sample. While steps 1-3 may be exploratory in nature, steps 4 and 5 should be conducted with a clear research question that cannot be answered using other research methods. Once a sample is deemed suitable for molecular analysis, careful consideration should be given to the sampling location and technique in order to preserve as much of the remaining bone structure as possible. CT scanning can be used to identify areas with sufficient bone density and well-preserved anatomical structures. When possible, researchers should prioritize areas that have already sustained damage or are less crucial for future morphological and histological studies (Rousaki et al., 2018).

**Figure 4.**
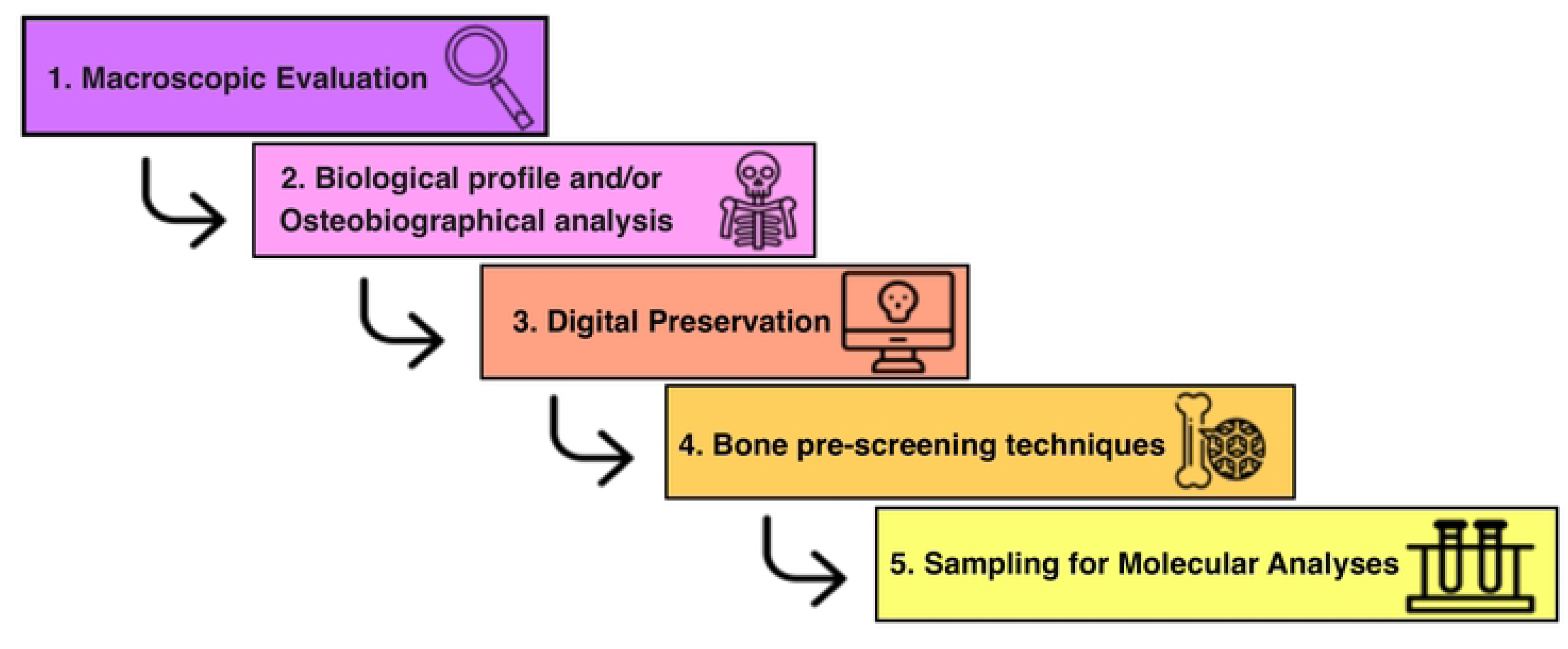
Workflow proposed for a sustainable and interdisciplinary approach to human evolution studies.

By following this workflow, molecular analyses can benefit from the access to relevant contextual information while optimizing time and financial resources. Identifying the anatomical elements present for each individual allows for the selection of the most suitable specimens for sampling. Assessing diagenetic features helps determine whether a sample, or specific portions of it, is viable for analysis. CT scanning also allows researchers to identify the densest areas within a specimen, which can guide the selection of regions most likely to yield well-preserved biomolecules (Alberti et al., 2018). Additionally, understanding an individual’s biological profile and lifestyle contributes to selecting the most appropriate samples for addressing specific research questions, while providing helpful contextual information for interpreting results from molecular analyses. Moreover, CT scanning, surface scanning, and photogrammetry methods ensure the digital preservation of samples, allowing researchers to revisit their original state before sampling and making the data accessible for future studies. X-ray fluorescence (XRF) analysis helps preserve valuable resources by preventing the unnecessary destruction of irreplaceable samples that may not meet the required compositional standards for further analysis. Finally, we recommend reducing the number of samples taken from the same individual for different analyses. Whenever feasible, tests such as radiocarbon dating, isotopes, and DNA should be performed using material from a single sampling event. CT scanning can aid this process by identifying optimal sampling areas and minimizing unnecessary damage.

## 6. Main challenges of imaging studies

We acknowledge that incorporating CT scanning into research workflows introduces additional steps into an already complex, expensive, and often time-consuming process for accessing and studying human skeletal remains. This additional step may significantly delay analyses, increase costs, and subsequent publication. To that end, here we outline the main challenges associated with this approach and propose potential solutions based on our collective experiences.

a. Challenges in accessing imaging equipment: Access to imaging equipment varies widely across the globe. In the most favorable scenarios, some institutions housing skeletal series and collections have their own µCT scanners, along with trained personnel to coordinate the logistics and conduct the scanning process (e.g., the Natural History Museum in Vienna, Austria). In contrast, other institutions are in remote areas where even medical CT scanners are unavailable within thousands of kilometers. Similar disparities exist for surface scanning and XRF, both of which depend on financial resources and trained personnel. To mitigate these challenges, researchers should allocate sufficient time to identify the most suitable conditions for imaging and collaborate with institutions that can provide access to the best available equipment and expertise. Establishing partnerships early in the research process can help secure the necessary resources and streamline logistical planning.
b. Financial constraints and potential solutions: The cost of CT scanning poses another significant challenge. With a global average of approximately $100 per scan, some research projects have sufficient funding to include CT imaging for multiple skeletons, while others have limited expenses or have not planned to allocate funding for CT scanning. Based on our experience working in countries from the Global South, alternative solutions can be explored when budgets are limited. One approach is to establish agreements with public medical institutions, universities or faculties (engineering, medical, chemical, veterinary) or collaborate with private medical facilities, which may provide access to CT scanners at reduced or no cost to the researcher.
c. Choosing between medical and micro-CT scanners for skeletal imaging: While medical CT scanners are more accessible, most times cheaper, and allow for scanning multiple bones at once, their resolution is considerably lower than that of micro-CT (µCT) scanners. As such, medical CT is better suited for larger anatomical structures (e.g., long bones, pelvis) and not recommended for skulls, where small structures like the inner and middle ear require higher resolution for clear visualization. However, in some cases, medical CT scanners may be the only available option. In such instances, we recommend using the highest possible spatial resolution (i.e., matrix), the smallest slice thickness, and selecting a scanning protocol that optimizes bone contrast.
d. Logistical considerations for sample transport: When imaging equipment is unavailable at the institution housing the human remains, researchers should arrange for sample transportation to a facility with CT scanning capabilities. Transporting the relevant skeletal samples requires meticulous packaging to prevent damage. We recommend using non-organic materials such as polyethylene foam, aluminum, bubble wrap, and sturdy plastic containers to ensure safe transit. Samples can be CT scanned in the same containers where they are stored (bags, boxes). In contrast, photogrammetry and surface scanning present fewer logistical challenges, as many surface scanners today are portable, and the work can typically be performed *in situ*. These methods primarily require access to electrical outlets, scanning equipment and software for image processing, making them more feasible in resource-limited settings. Photogrammetry and surface scanning require specific precautions to prevent contamination of samples. This includes stabilizing samples on a clean, sterilized surface, the use of gloves and masks to avoid direct contact, ensuring that tripods, cameras, and scanners are cleaned before and after use, and applying non-invasive stabilization methods that do not introduce foreign substances. Additionally, all work should be carried out in controlled environments where dust, skin particles, or chemical residues are minimized, and proper documentation of each step is critical to guarantee sample integrity and reproducibility of results.
e. Concerns about X-ray exposure and DNA preservation: Our results presented in Section 4 indicate that, when appropriate protocols are followed, conventional μCT scanning—and, in most cases, standard-dose clinical medical CT—generally do not reach radiation dose levels considered harmful to aDNA (Hall et al., 2015; Immel et al., 2016). In contrast, other X-ray imaging techniques such as Synchrotron scanning can exceed these thresholds and therefore require particularly stringent precautions (Immel et al., 2016). Under suitable conditions, DNA degradation does not surpass the natural rate of temporal decay, supporting the safe use of CT imaging in bioarchaeological research, provided that proper scanning protocols are applied.
f. The value of a collaborative approach: A collaborative approach that brings together morphologists, bioarchaeologists, archaeologists, curators, and paleogeneticists can significantly enhance sample selection by ensuring that research questions are matched with appropriate and sustainable analytical methods. Such collaboration aids in identifying well-preserved and complete specimens, thereby increasing the efficiency of molecular analyses and enabling more robust interpretation of integrated datasets. As shown in Section 5, our proposed workflow begins with an assessment of preservation, followed by osteobiographical analysis, selection for CT scanning, and ultimately radiocarbon dating and ancient DNA analysis.

## 7. Final considerations

As a multidisciplinary group of scientists specializing in morphology, paleogenetics, genomics, archaeology, and isotope analysis, we advocate for a responsible and sustainable approach to research, particularly when working with irreplaceable human remains. The petrous portion of the temporal bone has undeniably contributed to significant advancements in human evolutionary studies. As its demand continues to rise, however, it is critical to establish research protocols that prioritize conservation and efficiency without compromising scientific progress (Prendergast & Sawchuk, 2018; Squires & García-Mancusso, 2021; Cribellier & Chaillou, 2024; Källén et al., 2024; Menéndez et al., 2025).

The growing ethical awareness surrounding research on ancient human remains has prompted a welcome shift toward more careful stewardship and restricted access to collections. In this context, we recommend to prioritize responsible management, maximize the research potential of available materials, and promote transparency and data sharing to ensure that each study contributes meaningfully to broader scientific and ethical objectives (Ávila-Arcos et al., 2022; Prendergast & Sawchuk, 2018; Squires & García-Mancuso, 2021). These ethical imperatives demand methodological approaches that are not only minimally invasive, but also capable of generating the greatest possible amount of information across disciplines. With recent advances in biomolecular recovery techniques—such as the extraction of aDNA from buffer solutions applied to artifacts (Essel et al., 2023)—it is crucial to fully explore such alternatives before resorting to direct sampling of key structures like the petrous portion of the temporal bone, whose preservation is vital for both future research and heritage conservation.

A sustainable workflow ensures that valuable biological information to multiple disciplines is not lost due to excessive destructive sampling that only favours paleogenomic analyses. By integrating macroscopic evaluation, biological profiling, osteobiographical analysis, digital preservation, bone pre-screening techniques, and sampling for molecular analyses, researchers can maximize the analytical potential of each sample. Furthermore, promoting interdisciplinary collaboration among geneticists, morphologists, bioarchaeologists, archaeologists, and curators will facilitate data sharing, reduce redundancy in sampling efforts, and support the construction of more holistic understandings of past human lives (Menéndez et al., 2022).

Unequal access to imaging technologies and funding disparities remain major geopolitical challenges in implementing these protocols (Yáñez et al., 2023). Collaborative efforts between research institutions and public or private medical facilities can help bridge these gaps, allowing for broader access to imaging technologies such as CT scanning, surface scanning, and photogrammetry. Epistemic challenges—such as how to address inconsistencies between molecular and anatomical data, reconcile differences in timescales, and navigate hierarchies of evidence—remain to be thoroughly examined and integrated into comprehensive explanatory frameworks (Menéndez & Yañez, 2025). Furthermore, to foster future collaborations and ensure greater independence, researchers should plan for technology transfer and capacity building in regions lacking the necessary expertise and infrastructure, thereby helping to reduce asymmetries between researchers in the Global South and those in the Global North.

Ultimately, adopting a thoughtful scientific approach—one that emphasizes careful, methodical, and ethical research—ensures that valuable skeletal remains continue to inform multiple lines of inquiry while also being preserved for future generations (Prendergast & Sawchuk, 2018; Squires et al., 2019; Ávila-Arcos et al., 2022; Yáñez et al., 2023). The petrous portion is an invaluable resource, and its use in research should be guided by principles of sustainability, stewardship, and scientific responsibility. By refining research practices and fostering interdisciplinary collaboration, the field of biological anthropology can continue to produce meaningful insights while preserving the integrity of the subjects it studies.

Overall, our results demonstrate that µCT scanning does not compromise aDNA preservation. Building on this, we outline a sustainable workflow that integrates imaging, osteobiography, and compositional pre-screening before molecular sampling. This strategy promotes responsible stewardship of skeletal collections and fosters interdisciplinary collaboration, ensuring that the maximum information is obtained from each sample.

## Acknowledgments

We would like to thank Malena Vázquez and Julio Avalos from the Registro Nacional de Yacimientos Colecciones y Objetos Arqueológicos (RENYCOA) of Argentina for their assistance in the exportation of the samples studied. We would also like to thank the staff of the Faculty of Medicine, Paris Diderot University, for their assistance and for providing access to the scanning facilities. Finally, we would also like to thank the HPC Core Facility of Institut Pasteur (IP) for their support for computational analyses and Marc Monot, Laurence Motreff and Florence Jagorel from the IP Biomics Platform (supported by France Génomique ANR-10-INBS-09-09 and IBISA) for their assistance in sample sequencing.

## Data Availability Statement

All data necessary to reproduce the statistical analyses presented in this study are available in the supplementary material.

## Ethics Approval Statement

The sampling of archaeological human remains was conducted under the necessary permits for archaeological research in Argentina. The process of sampling and exporting the materials for ancient DNA (aDNA) analysis involved two steps. First, the issuance of a permit by the relevant cultural heritage authorities of the provinces where the remains were excavated and housed. Second, an export permit granted by the National Registry of Archaeological Sites, Collections, and Objects (RENYCOA), which operates under national jurisdiction to safeguard Argentina’s archaeological heritage and is responsible for authorizing the study of archaeological remains in foreign laboratories. The RENYCOA export permits for the samples analyzed in this study are: EX-2021-01946268, EX-2021-01968244, EX-2021-126859221, EX-2021-17680741, EX-2021-18124171, and EX-2021-18137399.

## Conflict of Interest Disclosure

The authors declare no conflict of interest.

Supporting information captions

Table S1. Archaeological metadata and molecular metrics of the samples included in the analysis.

